# Hydraulic tradeoffs underlie enhanced performance of polyploid trees under soil water scarcity

**DOI:** 10.1101/2022.09.24.509308

**Authors:** JM Losada, N Blanco-Moure, A Fonollá, E Martínez-Ferrí, JI Hormaza

**Author notes:** Author for correspondence: *Juan M. Losada, E-mail:*.

## Abstract

- Polyploid trees are excellent candidates to reduce crop water footprint and mitigate the increasingly reduced availability of freshwater for irrigation in many regions of the world due to climate change. Yet, the relationships between aerial organ morpho-anatomy of woody polyploids with their functional hydraulics under water stress remain understudied.
- We evaluated growth-associated traits, aerial organ xylem anatomy, and physiological parameters of diploid, triploid, and tetraploid genotypes of the woody perennial genus *Annona* (Annonaceae), testing their performance under long-term soil water reduction.
- Polyploids displayed contrasting phenotypes, vigorous triploids and dwarf tetraploids, but consistently showed stomatal size-density trade-off. The vessel elements in aerial organs were ∼1.5 times wider in polyploids compared with diploids, but triploids displayed the lowest vessel density. Sap flow velocity, measured *in vivo* through a novel method, was 10-fold faster in flower carpels than in second leaf vein orders. Triploid leaves displayed the slowest velocity in the leaves but the fastest in the carpels. Plant hydraulic conductance was higher in well-irrigated diploids at the cost of consuming more belowground water, but diploids showed less tolerance than polyploids to soil water deficit.
- The phenotypic disparity of atemoya polyploids associates with contrasting leaf and stem xylem porosity traits that coordinate to regulate water balances between the trees and the belowground and aboveground environment. Polyploid trees displayed a better performance under soil water scarcity, opening the possibility for deeper research on the factors underlying this behaviour and use them for a more sustainable agricultural and forestry production.

## Introduction

Plant evolution and species diversification are tightly linked with whole genome duplication (WGD) events (van der Peer *et al*., 2021). WGD occurs by two main processes, either by unreduced gamete formation (autopolyploidy) generating genetically identical individuals with multiplied chromosome numbers, or following interspecific hybridization (allopoliploidy), generating transgressive phenotypes. Plants show a much higher rate of allopolyploidy than other living organisms (Mable *et al*., 2011), and the combination of hybridization and chromosome doubling generates unique phenotypes with some degree of plasticity (Barker *et al*., 2016), therefore expanding their adaptive potential to stressful environmental scenarios (Wei *et al*., 2019; Baniaga *et al*., 2020). Indeed, some polyploid plants have been shown as more efficient in colonizing habitats that can be hostile for their diploid relatives, such as soils with high salinity and/or low water availability (Te Beest *et al*., 2012; Decanter *et al*., 2020). Most studies in either auto- or allopolyploids derive from herbaceous species, because polyploid trees are rarely found in the wild (Meyers & Levin, 2006), largely limiting our understanding on the consequences of polyploidy in the phenology, phenotype, and physiology of woody species (Hao *et al*., 2013; Zhang *et al*., 2017). In this sense, tree germplasm collections constitute excellent reservoirs of polyploids, in many cases including novel genotypes produced after long-term breeding and selection. Those genotypes may provide essential information for functional anatomy studies (Barceló-Anguiano *et al*., 2021), thus becoming pivotal in selecting woody crop genotypes better adapted to restrictive scenarios resulting from global climate change.

Among the multiple consequences of climate change, loss of primary productivity at the global scale associates with an increased vapour pressure deficit (VPD) in the atmosphere, linked with soil water scarcity in many regions of the world (Yuan *et al*., 2019). The consequences in agro systems of the most vulnerable regions, such as the Mediterranean basin, are aggravated by the intensification of the use of non-renewable water resources (Asner *et al*., 2016), picturing future scenarios with more frequent and prolonged droughts that will jeopardize food security and sustainable practices in agriculture (Wheeler & von Braun, 2013). Prolonged reduction of available belowground water may lead to the formation of embolisms in the water conducting pipes of plants (i.e. xylem vessels), with negative consequences on hydraulic conductivity, eventually causing hydraulic failure (Broddrib *et al*., 2011), massive dehydration of tissues, and, finally, drought-induced tree mortality (Choat *et al*., 2018). Indeed, the effects of extreme drought events in the vascular systems of trees have received increased attention in the last decade, and have been mostly evaluated in stems (Zieminska *et al*., 2020) or leaves (Sorek *et al*., 2021), but more scarcely in the reproductive organs (Roddy *et al*., 2019). In contrast, less attention has been given to the effects of prolonged soil water deficit in the physiology of trees, which requires monitoring the hydraulic performance of plants under different soil water availability scenarios (Anderegg *et al*., 2014; Brodersen *et al*., 2019).

Understanding the reasons behind the physiological effects of water scarcity in trees requires a deep knowledge of the architecture of the vascular conduits at the intraspecific level, given that dimensional variations of vessels along with their distribution in the plant greatly affect hydraulic performance (Roth-Nebelsick *et al*., 2021). Increased diameters of the vascular conduits have been reported in a handful of polyploid woody species (Allario *et al*., 2013 Hao *et al*., 2013; Zhang *et al*., 2017; Barceló-Anguiano *et al*., 2021), showing that the geometrical patterns of the vasculature are under strong genetic control (Brodribb *et al*., 2020). The physiological responses of polyploid plants under drought were already predicted 40 years ago (Levin *et al*., 1983), and they were tested in annual crops (Tal & Gardi, 1976; Yang *et al*., 2014; Pei *et al*., 2019). However, the effect of ploidy on phenotype, and more specifically on the vascular anatomy, is still mainly unexplored, particularly in woody species (reviewed in Ruiz *et al*., 2020), and, to the best of our knowledge, controlled water treatment trials have not been performed before in conspecific trees with increased chromosome numbers.

In order to fill this gap, we analysed diploid, triploid, and tetraploid hybrids of *Annona cherimola* x *Annona squamosa* (also known as ’
satemoyas’) cultivated under field conditions for over a decade. These genotypes belong to the species-rich genus *Annona* (Annonaceae, Magnoliids), composed of semi deciduous fruit trees with economic importance in many countries with tropical and subtropical climates. The Mediterranean coastal region of southern Spain is the main commercial producer of *A. cherimola* worldwide but, in light of climate change scenarios, productivity is jeopardized by the restrictions in freshwater availability for irrigation and, thus, developing new genotypes with higher water use efficiency is compulsory. *Atemoyas* constitute not only an excellent system in which to test the effect of soil water scarcity in both the physiology and growth of tree crops, but also a pioneer effort in developing strategies to confront the harmful effects of climate change in one of the most susceptible regions worldwide. We pursued two major goals: 1) To compare anatomical, photosynthetic and hydraulic traits of allopolyploid trees with their diploid relatives. 2) To understand how the functional anatomy of the xylem impacts the physiology of polyploids under sustained soil water deficit.

## Results

### Aboveground measurements of adult plants and grafted *atemoya* polyploids

Field grown trees of *atemoyas* display long branches with alternate nodes (Fig. 1a). Each node contains an average of four buds, two vegetative and two reproductive, and, although the four nodes have sprouting potential, typically just one or two of them - either reproductive or vegetative-develops during the growing season (Fig. 1b). Adult plants of all three ploidies grown from seeds displayed a direct correlation between branch width and length (*r*^2^=0.80; *p* < 0.05; Fig. S1) and, although the total number of nodes per terminal branch was similar across ploidies, the shorter branches of tetraploid trees resulted in a significantly higher node density (Fig. 1c; *F*=6.5; *p* < 0.05), further reflected in a higher flower density compared with diploids and triploids (*F*=3.5; *p* < 0.001). Leaf mass per area scaled directly with ploidy (Fig. 1d; *F*=17.9; *p* < 0.05), but the water mass per area was significantly larger in tetraploids (*F*=27.4, *p* < 0.001). Flower length to width ratio displayed an inverse relationship with ploidy (Fig. 1d, *p* < 0.05).

**Fig. 1.**
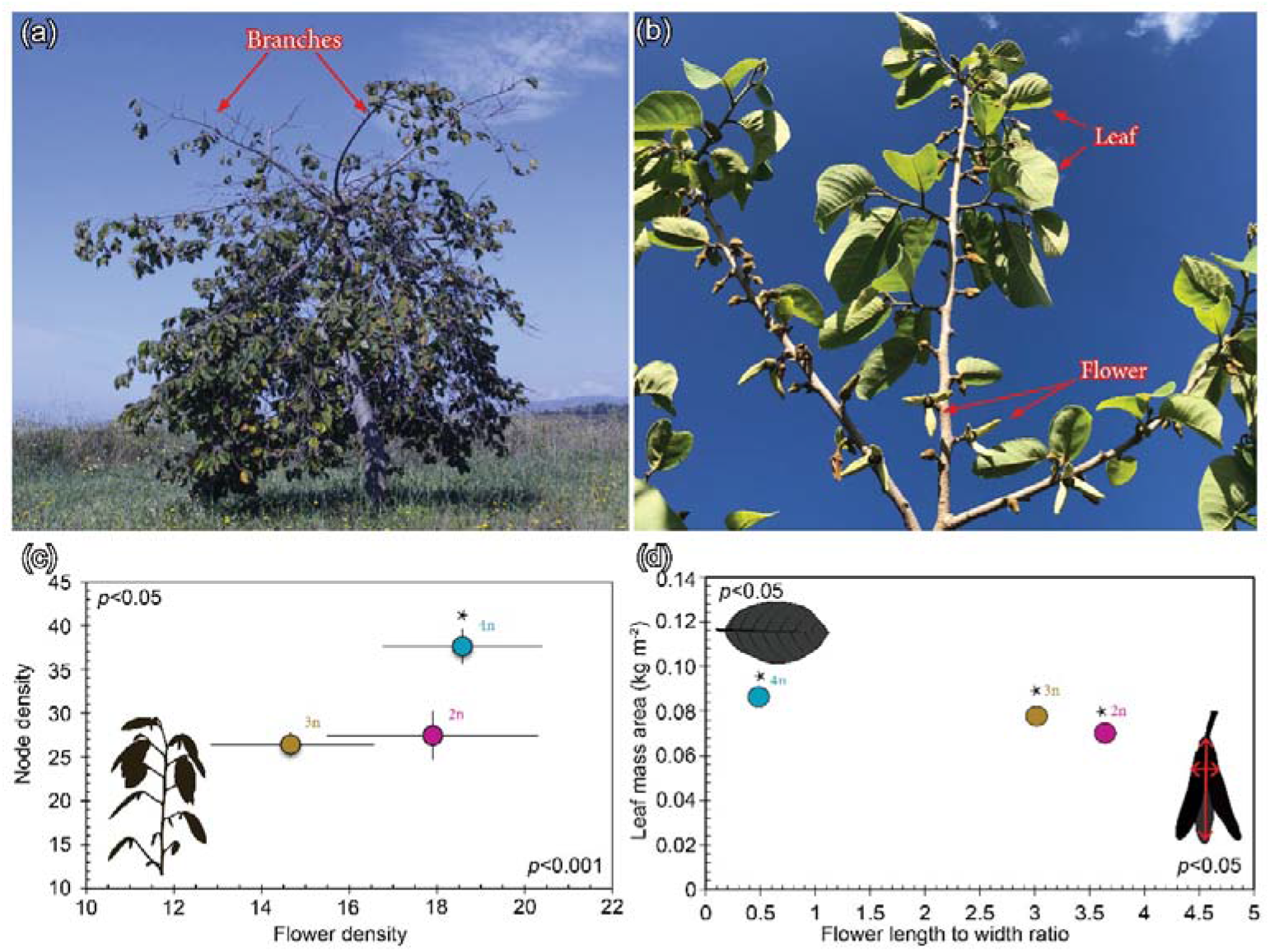
Vegetative and reproductive characterization of diploid (pink), triploid (brown), and tetraploid (blue) *atemoya* genotypes. **(a)** Adult tree of *Annona cherimola* in the field showing the terminal long branch pattern. **(b)** Detail of branches composed of alternate nodes from which leaves and flowers develop. **(c)** Relationship between node density and flower density per branch length (number m^-1^). **(d)** Leaf mass area (y axis) *versus* flower length to width ratio (x axis). Vertical and horizontal bars represent the standard error at *p*<0.05. Asterisk shows significant differences following a one-way ANOVA for each parameter represented in each axis at either *p*<0.001 or *p*<0.05 as noted.

The differences observed in aboveground organs of adult plants grown in the field were maintained in the grafted scions. Thus, the diameter of the branches was similar across ploidies (*F*=1.2; *p* = 0.30), but branch length was shorter in tetraploids and longer in triploids (*F*=4.9; *p =* 0.01) resulting in the highest and lowest node density respectively (*F*=2.9; *p* = 0.06). At the phenology level, diploid scions displayed a higher number of bursting nodes per branch in the beginning of the growing season, compared with the unburst nodes of triploids or tetraploids (April: *F*=4.5; *p* < 0.05), which delayed massive bud sprouting by 2-3 weeks. Comparing the number of leaves per branch revealed similar values for all ploidies in the beginning of the season, but tetraploids arrived at the end of the season with the lowest leaf number per branch (June: *F* = 4.4, *p* = 0.02, Fig. S2a). In an analogous fashion, foliar length revealed similar values across ploidies in the beginning of the season (April: *F*=0.02; *p* = 0.98; May: *F*=0.6; *p* = 0.56), but triploid leaves became significantly larger at the end of the expansion period (June: *F*=5.1; *p* < 0.05, Fig. S2b). Flowers were larger in polyploid scions, as observed in seedlings, but flower number varied across grafted scions, being higher in diploids (average 31.23±9.15SE flowers per tree), lower triploids (average 4.0±0.77SE flowers per tree), and almost no flowers developed in tetraploids.

### Anatomy of the aerial organs of *atemoya* polyploids

The stomatal complex of *atemoya* leaves displayed two elongated guard cells, surrounded by two parallel subsidiary cells, all embedded in irregularly shaped epidermal cells in the abaxial side of leaves (Fig. 2a-c). Cell size of the stomatal complex scaled directly with ploidy whereas stomatal density scaled inversely (Fig. 2d). These quantifications evidenced a correlation (*r*^2^=0.99) between the larger leaves of triploids and a significantly smaller (*F*=13.2; *p* < 0.05) stomatal index (SI: 19.2±0.41SE), whereas both diploids and tetraploids have comparable leaf areas and SI (22.5±0.52SE diploids, 22.2±0.58SE tetraploids).

**Fig. 2.**
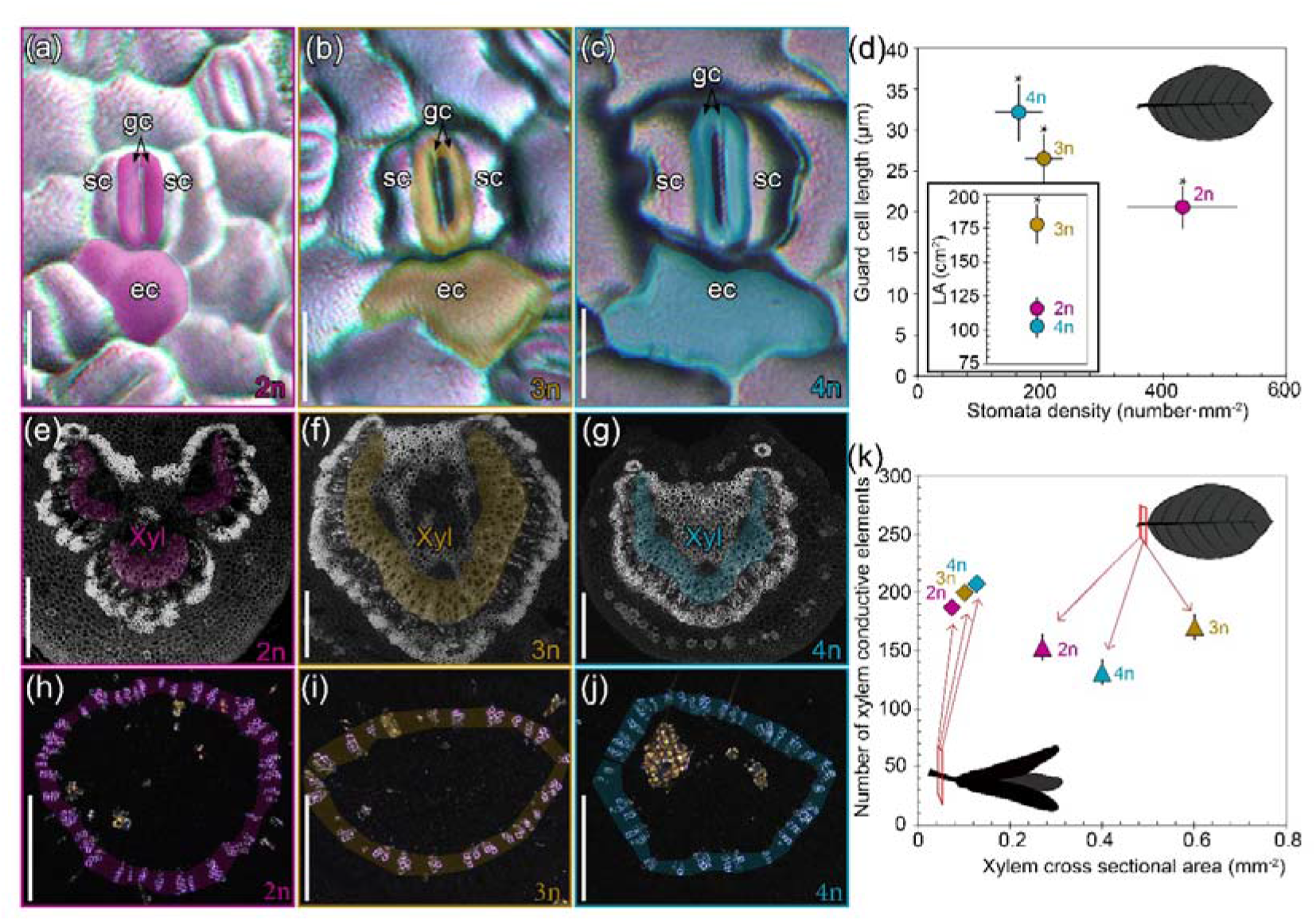
Anatomy of aerial organs of diploid (pink), triploid (brown), and tetraploid (blue) *atemoya* plants. **(a-c)** Stomata and abaxial leaf cells of the different ploidies in epidermal peels with polarized light. **(d)** Guard cell length and stomata density. Inset shows leaf areas. Asterisks display significant differences following one-way ANOVA at *p*<0.05, bars represent standard deviation **(e-g)** Cross sections of the petioles stained with calcofluor white and imaged with fluorescence, with colored areas emphasizing the xylem. **(h-j)** Cross sections of flower pedicels showing the tracheids of the xylem (colored areas) in semi thin sections with polarized light. **(k)** Relationship between number of conductive elements per section and total xylem area in the flower pedicels (ribbons), and in the leaf petioles (triangles) for each ploidy; bars represent standard error at *p*<0.05. Ec, epidermal cell; gc, guard cell; phl, phloem; sc, subsidiary cell; xyl, xylem. A-C scale bars: 20 μm; E-G, H-J scale bars, 500μm.

Cross-sections of leaf petioles exhibited a continuous pericyclic fiber layer surrounding an adaxial xylem populated with vessels (Fig. 2e-g). Both the average total vessel number per petiole (*F*=6.9; *p* = 0.002) and the xylem cross sectional area were significantly larger in triploids (*F*=12.4; *p* = 0.001) than in the other two ploidies (no significant differences were observed between diploids and tetraploids). As a result, on the basis of the total xylem area, triploids and tetraploids exhibited the lowest vessel density (283 vessels mm^-2^ for triploids and 328 vessels mm^-2^ for tetraploids) compared with the higher vessel density of the diploids (average 564 vessels mm^-2^).

The vasculature of flower pedicels at the anthesis stage was mainly composed of tracheids (Fig. 2i-k). Both total tracheid number (*F*=0.27, *p* = 0.76), and xylem cross sectional area (*F*=8.6, *p* = 0.01) scaled with ploidy, resulting in an inverse escalation of ploidy and tracheid density (diploids 2,510 tracheids mm^-2^; triploids 1,960 tracheids mm^-2^; tetraploids 1,620 tracheids mm^-2^).

The xylem of stems is diffuse-porous with interspersed ray parenchyma (Fig. 3a). The surrounding phloem is composed of concentric axial areas that decrease in size from the cambium to the cortex. While thinner stems up to 6mm diameter displayed similar xylem cross sectional areas and vessel element distribution across ploidies, in wider stems (9mm diameter), triploids displayed the lowest total vessel number per cross section (*F*=4.86, *p* = 0.03), and, therefore, the lowest vessel density of all ploidies (Fig. 3b-d).

**Fig. 3.**
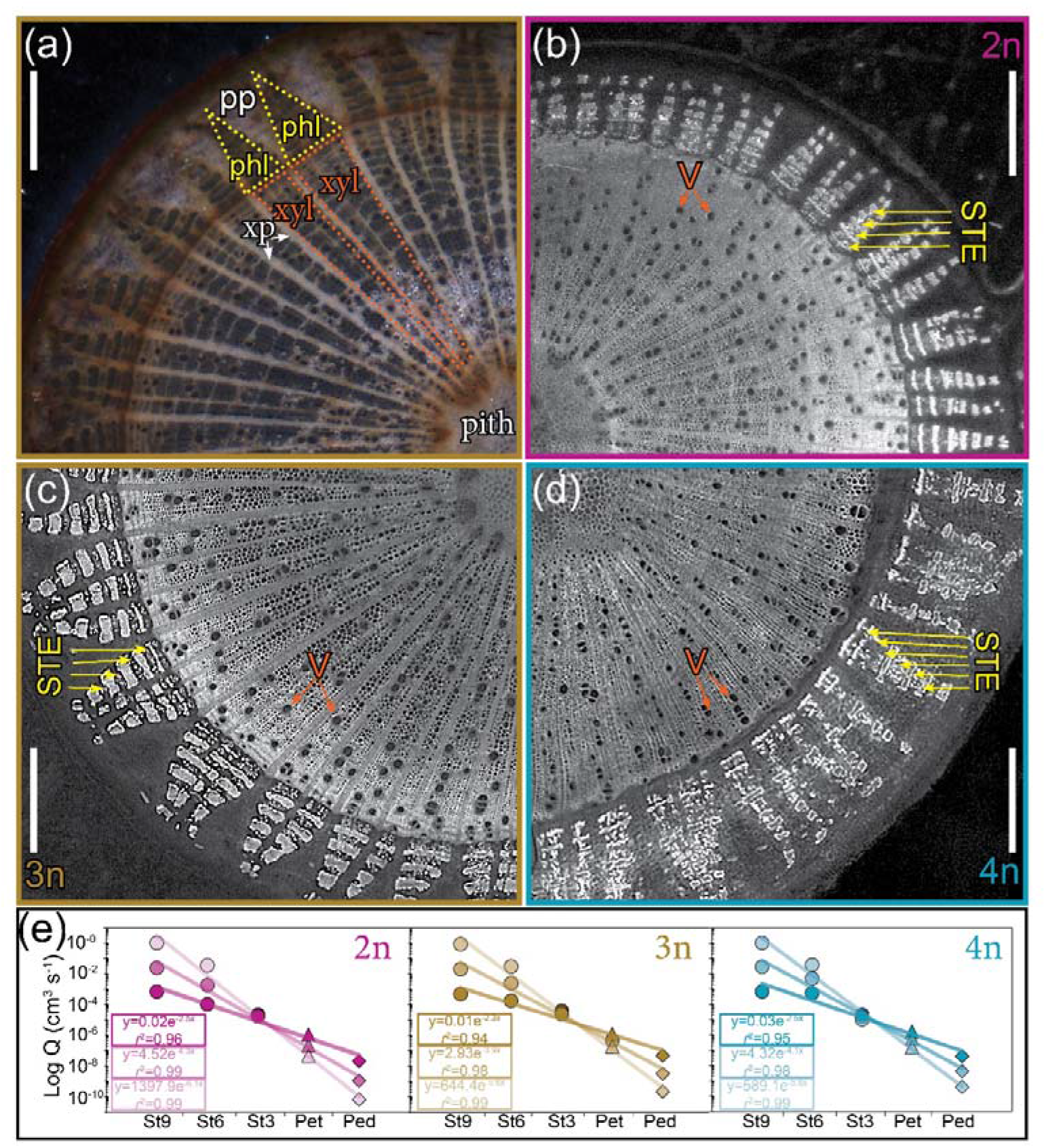
Anatomy of the stem and xylem sap flow rates in diploid (pink), triploid (brown), and tetraploid (blue) *atemoya* genotypes. **(a)** General anatomy of the stem of *atemoyas* in 9 mm cross sections, showing a central pith, xylem areas (orange dots), continued by axial areas of the phloem (yellow dots); both xylem and phloem are interspersed by ray parenchyma. **(b)** Cross section of a 9 mm diameter diploid stem stained with aniline blue that emphasizes the sieve tubes of the phloem (yellow arrows), and the xylem vessels (orange arrows). **(c)** Cross section of a 9 mm diameter triploid stem. **(d)** Cross section of a 9 mm diameter tetraploid stem. **(e)** Calculated xylem flow rates (Log Q) over 1m along the atemoya stems of decreasing diameters (circles), petioles (triangles), and flower pedicels (ribbons), based on anatomical and water potential measurements, depicting exponential decrease from the base to the apical parts of the trees; the degree of transparency represents less water in the soil. Ped, pedicel; pet, petiole; phl, phloem; pp, phloem parenchyma; St3, stem diameter 3mm; St6, stem 6mm diameter; St9, stem 9mm diameter; STE, sieve tube elements; V, vessels; xp, xylem parenchyma; xyl, xylem. Scale bars, 1mm.

The predicted maximum flow rate through the xylem revealed an inverse exponential escalation from the thicker to the thinner stems and to the leaf petioles/flower pedicels (Fig. 3e). Comparison of the slopes under gradual soil water reduction reveals sharper slopes with less water in the soil in all ploidies, but most especially in diploids, doubling the value of the polyploids.

### *In vivo* flow velocity through the xylem in leaves and flowers of *atemoyas* with different ploidies

*In vivo* velocity of dye perfusion in leaves displayed movement of the dye through the leaf veins of increasing orders (from the major vein to the minor veins (Fig. 4a-d, Video S2), and dissipation through the stomatal cell walls (Video S1), confirming the xylem pathway. Our generalized linear model with the aggregated data for veins of different calibres revealed that the differences observed in bulk flow velocities were explained by ploidy, by vein thickness, and by their intersection (*r*^2^ = 0.94; *p* < 0.001). To homogenize comparisons across ploidies, we selected veins with a thickness between 20 and 30μm, given that vigorous triploid leaves rarely contained thinner veins. The secondary veins of triploid leaves displayed significantly lower speeds (5.9±0.5SE μm s^-1^; *F*=15.6, *p* < 0.001) compared to the higher speed in diploids (17.3±3.0 μm s^-1^) and tetraploids (20.9±5.4 μm s^-1^), which showed no significant differences between them (Figure 4i).

**Fig. 4.**
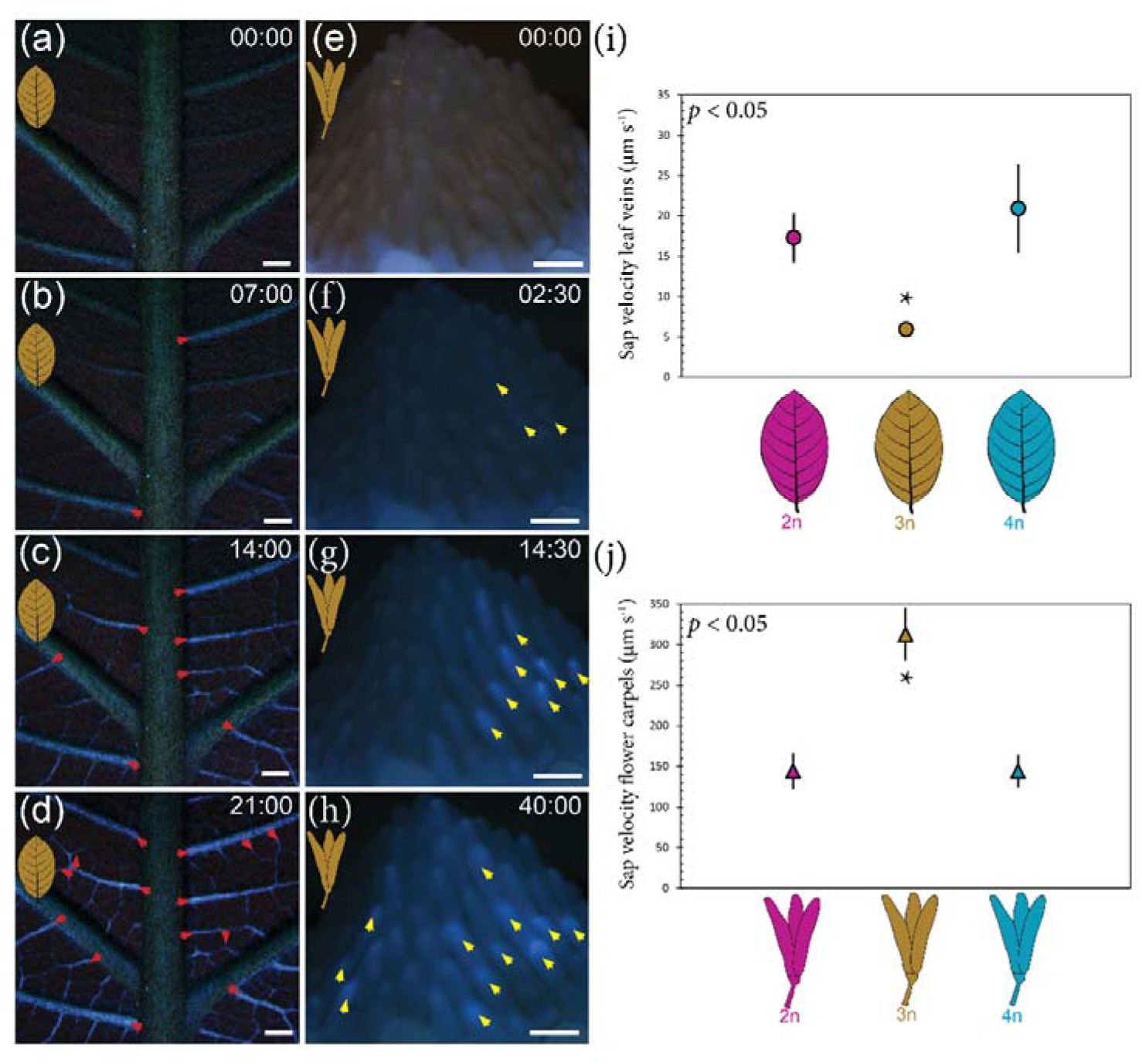
Measurement of *in vivo* xylem sap speed in leaf veins and flower carpels of *atemoya* polyploids. **(a-d)** Fluorescence track of esculin perfused through the xylem in a triploid leaf, showing the hierarchical stain through veins of increasing order (red arrowheads). **(e-h)** Time lapse of esculin transport through the individual flower carpels of triploid flowers (yellow arrowheads). **(i)** Speed of esculin directional transport through the secondary veins of 20-30μm thickness comparing diploid (pink), triploid (brown), and tetraploid (blue) leaves. **(j)** Esculin accumulation in individual carpels of diploid (pink), triploid (brown), and tetraploid (blue) flowers. Stars display significant differences in velocity across ploidies at *p*<0.05 **(a-d)** scale bars, 1000μm; **(e-h)** scale bars, 500μm.

Dye perfused in the flower pedicels (more than 2 cm away from the carpels) was gradually accumulated in the carpels (Fig. 4e-h), and dissipated afterwards through the stigmatic surfaces (Video S3). Comparisons of the slopes of pixel accumulation curves with time across ploidies revealed that dye velocity in carpels was tenfold faster than in the secondary veins (Fig. 4j). The carpels of triploid flowers showed the highest velocity (329.5±32.5SE μm s^-^1; *F*=13.9, *p* < 0.001) compared to the tetraploids (159.4±20.3SE μm s^-1^) and diploids (156.2±22.1SE μm s^-1^), without significant differences between the latter (Fig. 4k).

### Soil water balances and plant growth traits in grafted *atemoya* polyploids under different irrigation intensities

Both the relative humidity and the temperature in the greenhouse were quite stable during most of the year, except during summer when both increased in 2019 and 2020 (Fig. S3). The soil matric potential under the three irrigation regimes remained close to zero throughout the fall and winter seasons (note that flowers do not self-pollinate and thus, fructification was avoided, suppressing fruit sink strength Fig. 5), reflecting a low water consumption by plants in this period. However, soil matric potential dropped concomitantly with leaf emergence, and reached a minimum at the fully expanded leaf stage (Fig. 5a), with more negative values under low (4L/week) and medium (8L/week) irrigation treatments (values corrected for soil evaporation). Under good irrigation (16L/week), soil water potentials decreased considerably only in the soils of pots with diploid trees. We applied a model to the aggregated data of stem girth increase over a year, which revealed that variation was partially explained by ploidy (*F*=3.9, *p* = 0.08), by year (*F*=179.2, *p* = 0.01) and irrigation treatment (*F*=15.3, *p* = 0.04). Thus, well-irrigated diploid plants increased stem girth by four times over a year, compared to the three times increase in well-irrigated polyploids (Fig. 5b). Comparisons within ploidies showed that variation between years was significant in well-irrigated diploid (*F*=21.8, *p* = 0.01) and triploid (*F*=15.5, *p* = 0.04) trees, but no differences among the different irrigation treatments were observed in tetraploids (*F*=1.5, *p* = 0.28). In addition to the stem, irrigation affected the total leaf number per branch depending on the ploidy: diploids scaled leaf number with water applied to the soil (Fig. S4), but tetraploids displayed the inverse effect; triploids showed the maximum leaf number per branch in the medium irrigation treatment.

**Fig. 5.**
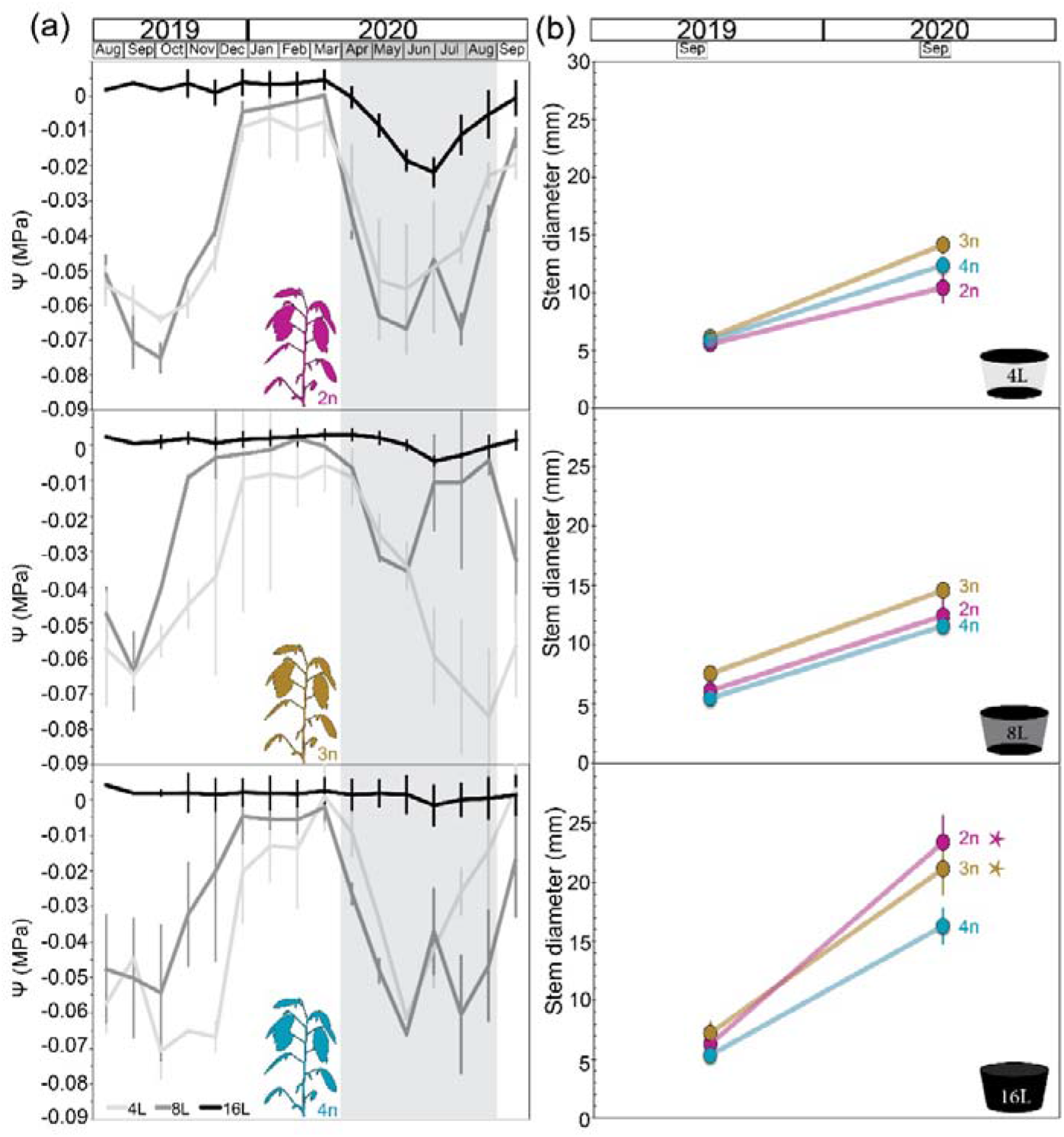
Belowground and aboveground variables over a two-year irrigation experiment in *atemoyas*. **(a)** Soil matric potential of the diploid (top plot), triploid (mid plot), and tetraploid (bottom plot) genotypes under three irrigation regimes per week: four liters (light grey), eight liters (dark grey), and sixteen liters (black). The shaded area represents spring-summer period of 2020, which corresponds with the growth season. Bars show the standard error of the monthly averages (corrected with the negative controls) at *p*<0.05. **(b)** Branch diameter during 2019 and 2020 under the three irrigations described in diploid (pink), triploid (brown), and tetraploid (blue) genotypes. Bars represent standard error and star significant differences within the same ploidy across treatments, both at *p*<0.05.

### Physiology of grafted *atemoya* polyploids under different irrigation intensities

A generalized linear model revealed a significant effect of the irrigation intensity in most gas exchange variables (Table SI), such as assimilation rate (A), stomatal conductance (Gs), internal CO_2_ (C_i_) and evapotranspiration (E). Additionally, assimilation rates and C_i_ varied across ploidies (Fig. 6): assimilation rates were low under the low irrigation treatment and increased linearly in both diploids and triploids when more water was applied to the soil, whereas no significant differences were observed in the tetraploids (Fig. 6a). Although assimilation rates were similar in all ploidies under good irrigation, diploids had higher stomatal conductance and, thus, showed the lowest intrinsic water use efficiency (Fig. S5, Fig. 6b). In line with this, diploids exhibited low evapotranspiration values under water scarcity, but higher evapotranspiration when water in the soil was increased. Values in the tetraploids remained similar under the three irrigation intensities (Fig. 6c), resulting in a higher instantaneous water use efficiency (Fig. 6d). Despite the similar stomatal conductance of all ploidies under medium and high irrigation intensities (Fig S5), diploids showed the highest leaf internal CO_2_ concentration (Fig. 6e, f).

**Fig. 6.**
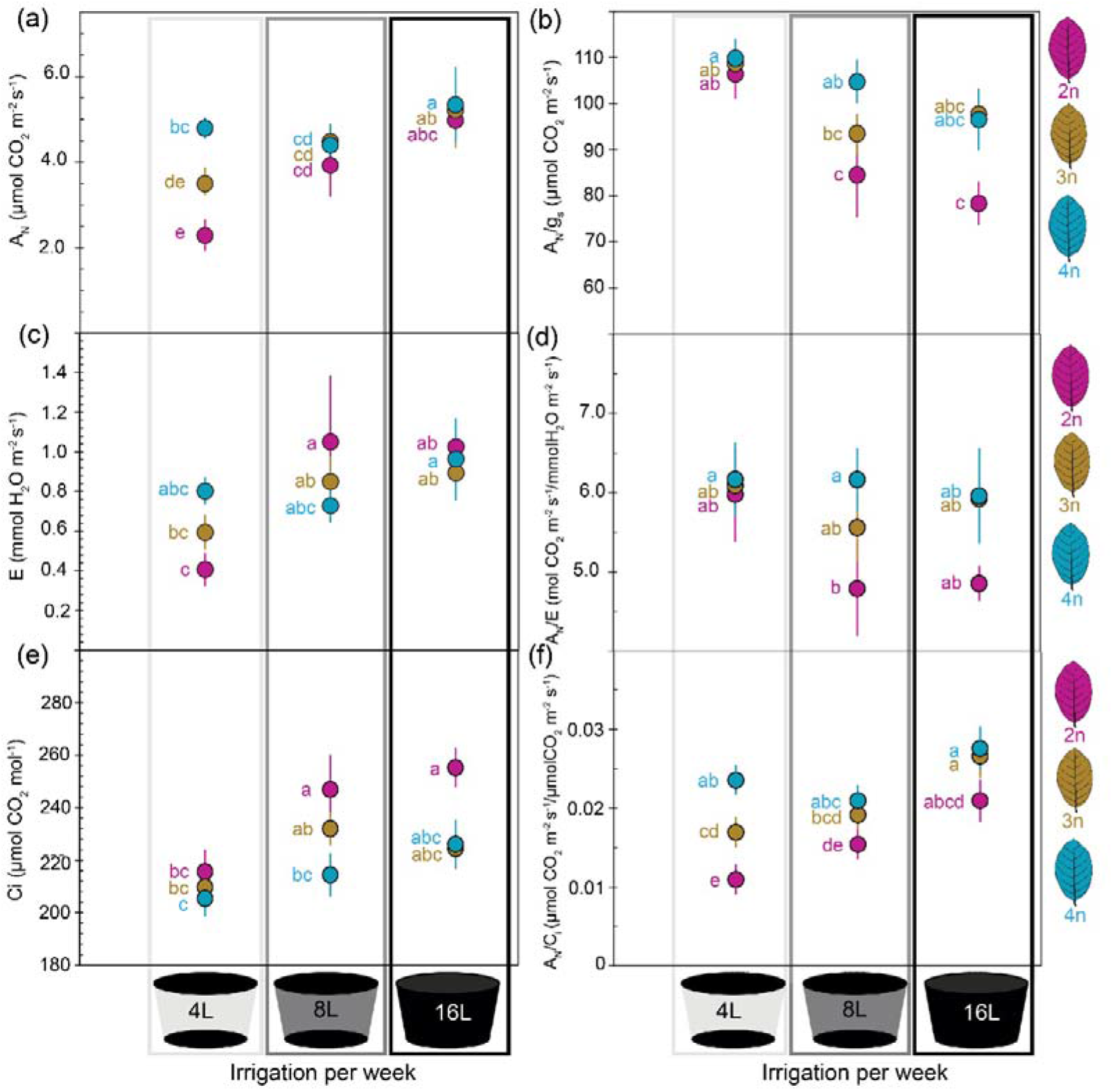
Physiology of *atemoya* diploid (pink), triploid (brown), and tetraploid (blue) genotypes under three irrigation regimes. **(a)** Assimilation rates of tetraploids remained stable, whereas a direct correlation between assimilation rates and irrigation was observed for diploids and triploids. **(b)** The relationship between carbon assimilation and stomatal conductance was lowest in diploids under good irrigation. **(c)** Evapotranspiration rates was maximal under good irrigation for all ploidies. **(d)** Water use efficiency was minimal for diploids under good irrigation. **(e)** Internal CO_2_ concentration was higher in diploids. **(f)** The relationship between assimilation and internal CO_2_ was minimal for tetraploids under all irrigations. Bars represent standard error at *p* < 0.05. Irrigation regimes per week are represented in different grey scales, from the lowest (4L, light grey), medium (8L, dark grey), and high (16L, black). Letters represent significant differences after a two way ANOVA at *p* < 0.05.

Field-grown plants, which were irrigated plentifully, showed no differences in neither gas exchange variables (Table SI) nor leaf water potentials (Ψ_l_) compared with well-watered grafted plants in the greenhouse (Fig. 7a). Although values of leaf water potential were high for all cytotypes, they displayed significant differences when comparing predawn (between 0 and -0.3MPa) and midday (between -0.6 and -0.9MPa) potentials (*F*=150.5; *p* < 0.001). Thus, trees with low water in the soil reduced the Ψ_l_ difference between midday and predawn. Integration of Ψ_l_ at midday with soil matric potential (Ψ_S_), and plant evapotranspiration (E) pictured that plant hydraulic conductance (K_plant_) was higher all ploidies under medium and high soil water scenarios, but differences were observed for K_plant_ when ploidy and irrigation were taken together (F=185.4, *p* < 0.001), especially when comparing extreme irrigation treatments, depicting a sharper reduction of K_plant_ in diploids (Fig. 7b).

**Fig. 7.**
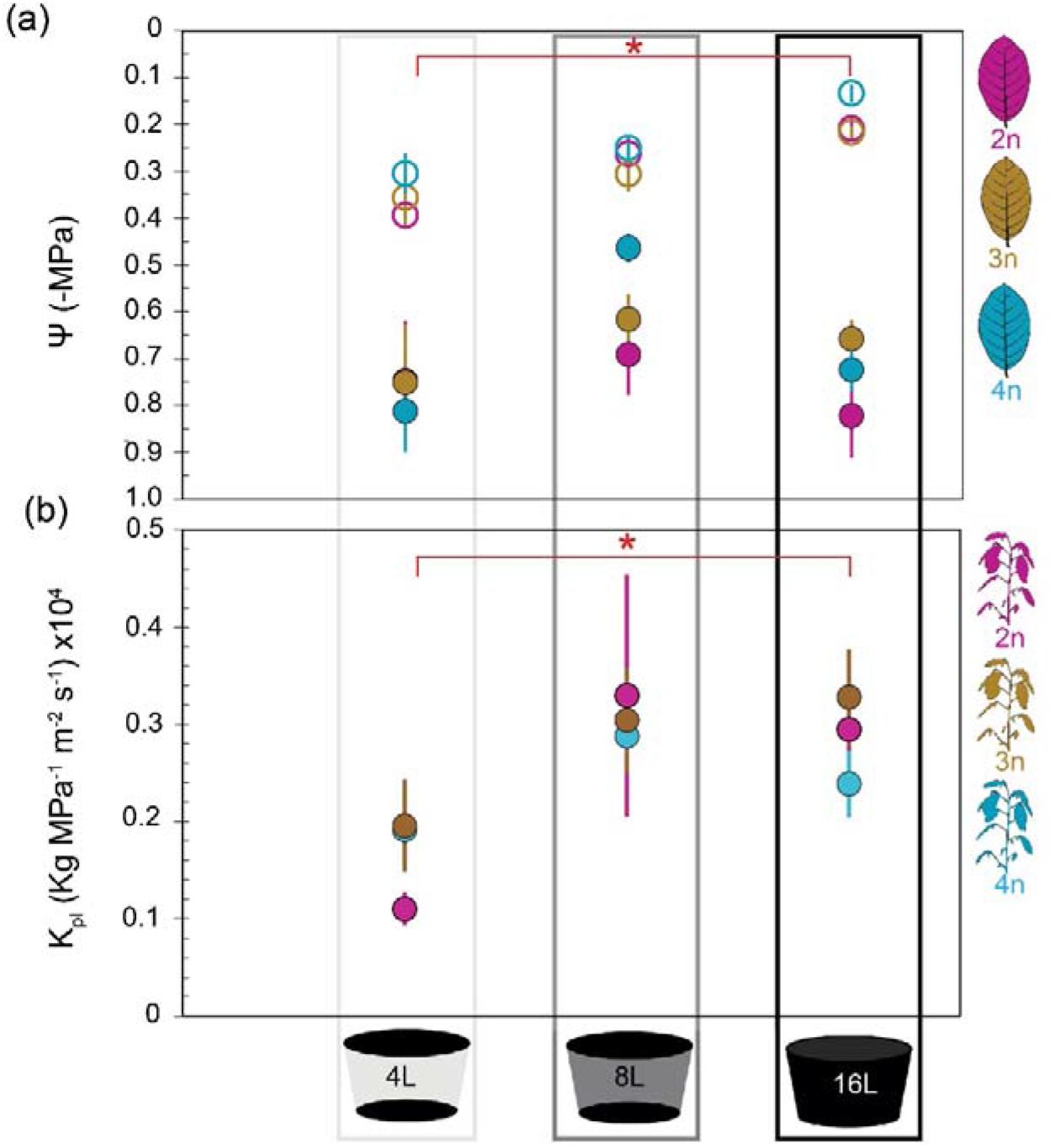
Plant hydraulic conductance as a function of ploidy and irrigation. **(a)** Leaf water potential (Ψ) revealed that predawn Ψ (empty circles) are significantly more negative in water stressed plants (4L) than in well-irrigated (16L) plants. In contrast, midday Ψ (filled circles) showed no differences between ploidies or irrigations intensities. **(b)** Plant hydraulic conductance from soil to leaf displayed overall higher values for triploids and diploids under high and medium irrigation intensities, but plant hydraulic conductance decreased sharply in diploids under poor irrigation, compared with the less affected polyploids. Bars represent the standard error, and the asterisk shows significant differences between extreme treatments, both at *p* < 0.05

## DISCUSSION

By combining the characterization of functional anatomy with physiological parameters of trees in both open field and controlled greenhouse conditions, we derive empirical evidence of the anatomical variation underlying the physiological advantage of polyploid trees under maintained soil water scarcity, and provide with a new tool to evaluate sap flow *in vivo*.

### Phenotypic diversity of polyploids: from whole plants to cells

Our observations revealed that flower and leaf emergence occur earlier in atemoya diploids than in polyploids. Polyploids have been previously stated as more susceptible to modifications of the reproductive function (Martin & Husband, 2012), a feature that is critical for fitness in natural populations (Gaudinier and Blackman, 2020). In the case of leaf phenologies, polyploids could also show adaptive functions, such as the ability to confront the erratic climate events during the growing season, thus with implications on the survival and colonization of new habitats (Simón-Porcar *et al*., 2017). This is particularly relevant for cultivated species such as of the genus *Annona*, since fruit trees with asynchronous flowering times might be interesting to increase the temporal frame of the reproductive period, therefore increasing the opportunities to optimize a minimum fruit production under uncertain climate scenarios.

The question remains on whether phenology swifts could be associated with phenotypic variation. We report phenotypic disparity of conspecific allopolyploids, exhibited as dwarf-like tetraploids with smaller branches with higher node density, and vigorous triploids with larger branches, but lower node density. The phenotypic effects of ploidy changes at the whole plant have been reviewed for herbaceous and woody plants (Ruiz *et al*., 2020), and previous works in different genera, such as *Betula* (Mu *et al*., 2012), *Malus* (Ma *et al*., 2016), and *Salix* (Dudits *et al*., 2016), emphasized a correlation between the widespread dwarfism of autotetraploid trees and altered levels or plant growth regulators. Our results are in line with these observations, but further reveal enhanced vigor of the triploids, similar to previous reports in *Salix* (Serapiglia *et al*., 2015) and *Populus* (Greer *et al*., 2018). The molecular mechanisms underlying triploid vigor are not yet well understood, but increasing evidence points to a combination between heterosis and genomic dosage as the main factor influencing the higher vigor observed in some polyploids (Chen, 2010). Interestingly, although the higher vigor of triploids is related with larger leaves and flowers, tetraploids showed contrasting characters, such as larger flowers but similar leaf size compared to diploids. Previous works emphasized a direct scaling relationship between ploidy and leaf size (Dudits *et al*., 2016; Serapiglia *et al*., 2015), but this relationship is not linear, as observed in *Arabidopsis*, where the larger leaves of tetraploids contrasted with the smaller leaves of octoploids (Kondorosi *et al*., 2000), emphasizing the adaptive divergence that follows post polyploidization events (López-Jurado *et al*., 2022). Regardless of organ size, leaf mass area scaled with ploidy in atemoya, in line with previous observations in other woody plant genera such as *Lonicera* (Li *et al*., 2009), *Populus* (Zhang *et al*., 2019), or *Mangifera* (Barceló-Anguiano *et al*., 2021), a feature that has been previously associated with a higher deposition of different organic compounds in the cell walls (Corneillie *et al*., 2019). In fact, the correlations between ploidy and organ size are typically perceived from the perspective of a tradeoff between cell size and cell density. Our measurements in mature leaves confirmed that both guard cells of the stomata and abaxial epidermal cells increased in size with ploidy, whereas their density per leaf area decreased, consistent with previous observations in other polyploids (Schnitzler & Muller, 1998; Beaulieu *et al*., 2008; Schlaepfer *et al*., 2010; Roddy *et al*., 2020; Barceló-Anguiano *et al*., 2021). Stomatal size and distribution have important physiological implications (Bomblies, 2020), through the relation between water tension within the xylem and stomatal dynamics, therefore conditioning the hydraulics of the whole plant (Meinzer *et al*., 2009). Among the multiple factors influencing stomatal behaviour, larger guard cells of polyploids may condition their aperture dynamics, but their distribution in the leaf is also relevant. Thus, despite their higher cell sizes, triploids displayed a smaller stomatal index, a parameter that relates with the intensity of transpiration (Brodribb & McAdam, 2017). Although this has been largely studied in different plant species, the correlation between stomatal index and the distribution of the vascular conduits of the xylem has remained understudied, is spite of the fact that this interaction influences hydraulic performance at the whole plant level.

### Tradeoffs between vessel diameter and packing explain the variable hydraulic function of polyploids

Our evaluations of the xylem architecture in the aerial organs at different levels provides another evidence of the general increment of vessel diameter with ploidy in aboveground organs, as recently reported in *Mangifera indica* trees (Barceló-Anguiano *et al*., 2021). A positive correlation between vessel diameter and ploidy has been reported in different angiosperm species, including herbaceous [*Chamerion* (Maherali *et al*., 2009)], and woody [*Atriplex* (Hao *et al*., 2013), *Populus* (Zhang *et al*., 2019), *Malus* (De Baerdemaeker *et al*., 2018)] genera. Our comparisons showed no differences in vessel size between triploids and tetraploids, suggesting that increases in ploidy confront anatomical limitations of vessel width expansion. Larger vessel diameters associate with the enhanced hydraulic efficiency of polyploids, which at the same time conflicts with embolism vulnerability (Sperry *et al*., 2008), although the relationship between vessel diameter and vulnerability is still under debate (reviewed by Lens *et al*., 2022). Despite vessel size differences, triploids exhibited a lower xylem porosity in the leaves and stems, whereas diploids showed higher leaf and stem porosity; xylem of the tetraploids had an intermediate phenotype, less porous in the leaves, but relatively porous in the stem. Contrasting xylem patterns between leaves and stems in tetraploids might associate with a possible conflict between the hydraulic architecture and the transpiration capacity, meaning that, in order to keep xylem tensions comparable to other ploidies, they would need to open stomata more broadly, during a longer period of time, or both. When these anatomical features are combined with the measured water potentials of trees, the predicted maximum water flux according to the Darcy’
ss law (Leyton, 1975), revealed that tetraploids exhibit greater volumetric sap flow rates from the stems to the leaves, which correlates with their higher xylem porosity in stems, and the faster sap velocities under good irrigation. Differences among cytotypes became more obvious with water scarcity, revealing drops in the flux slope between extreme zones of the transport pathway (the base of the stems and the leaves/flowers), that are sharper for diploids, suggesting their higher susceptibility to increases in water potential differences between soil and leaves. Estimations of the sap flow rate following Darcy’s law ignore the contribution of water retention in other cells composing the xylem, such as fiber cells, abundant in the less porous but more robust triploids, but their water capacitance remains to be tested. One of the main inconveniences when predicting flow rates is the availability of a reliable method to evaluate sap velocity within the xylem *in vivo*. In order to overcome this, we developed a visual method that tracks xylem sap speed through the movement of a dye in leaves and flowers *in vivo*. The results showed that the bulk speed of the xylem sap in flowers was tenfold faster than in the leaves, and that triploids showed lower speeds in the leaves but higher speeds in the carpels compared with the other two ploidies. While the differences in velocity between vegetative and reproductive organs could be associated with the type of transport, linear in leaves and cumulative in the carpels, the contrasting pattern may also respond to different regulatory mechanisms underlying flow velocity in source and sink organs. In source organs, the velocity is mainly controlled by stomatal conductance, whereas in sink organs, the underlying forces are less understood (Sinha *et al*., 2022). Previous works evidenced that a variety of hydraulic traits governed hydration of flowers (Roddy *et al*., 2019). Within flowers of *Annona*, the tip of the carpels (stigmas) contain an extracellular secretion rich in sugars (Lora *et al*., 2010), and this osmotic potential may also play a role in the directional transport of water. It is worthy to re-emphasize that velocity patterns in triploid leaves were in line with the measured anatomical parameters (lower xylem leaf porosity), and the predicted sap flow rates, smaller than in diploids and tetraploids. Previous evaluations of the sap flow speed through the xylem used detached leaves (Moghimislam *et al*., 2009), and there was a lack of information on the process in flowers, thus hampering comparison among species. As a result, the novel method described in this work appears as an easy and attractive approach for *in vivo* measurements of xylem bulk flow rates with application in a wide range of species.

### Enhanced hydraulic conductance of atemoya polyploids under soil water scarcity

To understand how and to what extent anatomical differences between diploids and polyploids are related to physiological responses under water stress, greenhouse experiments simulating different soil water availability scenarios revealed significant patterns associated with ploidy and irrigation intensity. The higher irrigation intensities resulted in a linear increase in the number of leaves and stem diameter in diploids, but no significant changes were observed in stem width in tetraploids, that showed a striking decrease in the number of leaves when more water in the soil was available. Triploids reached similar stem diameters than diploids, but consuming less water from the soil. In fact, regardless of the similar CO_2_ assimilation rates across ploidies under good irrigation, diploids revealed a lower water use efficiency. Although diploids appeared to be successful under good irrigation conditions, under soil water scarcity conditions hydraulic variables pointed to diploids as more susceptible. Among the few previous evaluations of physiological performance of polyploids under water stress, the variable responses to abiotic stresses pointed to be quite species-specific (Oliveira *et al*., 2017; Liao *et al*., 2018; Xu et al., 2018; Mtileni et al., 2019). However, general patterns such as a general increase of the assimilation rate of tetraploids correlated with an overall higher water use efficiency (reviewed in Ruiz *et al*., 2020). Our results confirm this pattern, but, in addition, suggest that the physiological effects depend on the type of ploidy and the extent of irrigation. These features are important for adaptive traits of polyploids, which exhibit phenotypic plasticity (Hao et al., 2013; Zhang et al., 2017). Our study is pioneer for trees, but it is important to keep in mind the life history of the Annonaceae, plants adapted to tropical and subtropical environments with high water demands (Awachare *et al*., 2018). For example, the similar midday leaf water potential among ploidies might correlate with a high drought susceptibility that matches with other species grown in subtropical climates (McDowell & Allen, 2015). Predawn leaf water potential was more negative in poorly irrigated trees of all ploidies compared with fully irrigated trees, pointing to a higher tension in the xylem under water scarcity at night, when stomata are supposedly closed. *Annona* cytotypes may therefore ameliorate excessive water loss under drought scenarios by reducing the day to night xylem tension. However, the mechanisms underneath remain to be elucidated.

Whole plant hydraulic conductance of *Annona* cytotypes was low compared with observations in temperate tree species (Nardini & Salleo, 2000; Becker *et al*., 1999; Rodríguez-Gamir *et al*., 2016), perhaps reflecting their adaptation to higher environmental humidity conditions. However, even with high atmospheric humidity, our results reveal that plant hydraulic conductance dropped faster in diploids than in polyploids under poor irrigation intensity. These results strongly suggest that diploid genotypes of atemoya are more efficient in scenarios of water abundance, producing more aboveground organs than genotypes of higher ploidies, but at the cost of consuming more belowground water. However, their sensitivity to water stress, likely through a rapid closure of stomata, seems to be higher than that of their polyploid relatives. As a consequence, diploids may be more efficient during short periods of drought, but they will suffer at the long term under dry conditions due to the starvation of stored energy resources. On the other hand, polyploids cope with water deficit in the soil despite their contrasting architectures of vessel and stomatal densities, and, while these combinations confer adaptation to prolonged drought periods, survival might be compromised due to the loss of hydraulic function, consequence of, for example, embolism spread that may cause hydraulic failure (Choat et al., 2018; Mantova et al., 2021). Enhanced water use efficiency of polyploid trees pointed to their lower susceptibility to water scarcity scenarios (Barceló-Anguiano *et al*., 2021; De Baerdemaeker *et al*., 2018; Oliveira *et al*., 2017; Liao *et al*., 2018) and thus our work provides empirical evidence of a robust performance of polyploid trees under drought. Our results further point to the possibility of using polyploid rootstocks with enhanced ability to tolerate soil water scarcity combined with diploid varieties that have been selected over years to attenuate the demands of water in crops with high water consumption.

## Conclusions

Combined detailed anatomical evaluations with novel combinations of interploid grafting and controlled irrigation served to understand that allopolyploid scions of trees conserve their diverse phenology, phenotype, and hydraulic function, all resulting in an enhanced performance under low soil water availability. These results are novel from the evolutionary perspective of adaptation of plants to water stress and, therefore, open exciting opportunities for the use of polyploids as novel genotypes with increased drought resilience, either as fruit trees or as biomass accumulators.

## Methods

### Plant material

Adult eight-year-old atemoyas (*Annona cherimola* x *A. squamosa*) grown from seeds and maintained under field conditions and irrigated on demand with an average of 72L/week were used as reference for anatomical and physiological evaluations, as well as scion donors for grafting in order to perform the experiments in controlled conditions in the greenhouse. Those trees were the second generation of backcrosses between the F1 of an interspecific cross (*A. cherimola* x *A. squamosa*) with the maternal parent (*A. cherimola*). Young leaves of each of these trees were analyzed with a Cyflow® PA flow cytometer (Sysmex), using a sample with known diploid nuclei from *A. cherimola* (Dolezel *et al*., 1989), revealing the presence of several diploid individuals and a few triploids and tetraploids (Martín *et al*., 2019).

### Morphology of aerial organs in adult plants and grafted atemoya polyploids

To understand dimensional differences in vegetative and reproductive organs (traits associated with plant growth) of field-grown adult plants from different ploidies, we measured ten terminal branches per tree (n=30), recording length, width, and the density of nodes and number of flowers. Furthermore, leaf area was evaluated in ten fully expanded leaves per ploidy (n=30) and weighted with a precision scale before and after dehydration at 60^°^C, to calculate leaf mass per area (LMA, kg m^-2^), and the water mass per area (WMA, kg m^-2^), as to (SW-DW)/A, being SW, saturated weight (kg), DW, dry weight (kg), and A, area (m^-2^). In the flowers, length and width of the tepals protecting the reproductive structures were measured at the anthesis stage in ten flowers per ploidy (n=30) using a calliper.

### Leaf and stem anatomy of atemoya polyploids

Stomatal morphology was measured in five fully expanded leaves per ploidy by the nail polish printing profile method (Karabourniotis *et al*., 2001), and images for measurements of these epidermal impressions, that included guard cells, subsidiary cells and epidermal cells, were obtained with a Leica DM LB2 microscope (n=150 stomata). Stomatal index (SI) was calculated as [SD/(SD+ED)] x 100, where SD is the number of stomata measured per unit area, and ED, the number of epidermal cells per unit area.

In order to evaluate the vessel diameter, vessel element density and the porosity of the xylem (number of conducting elements per mm^-2^, and total xylem area, respectively) in different axial positions of the stem, we measured three consecutive cross sections from at least 3 stems per ploidy and two diameter ranges (2-6mm and 9-11mm), from 5 flower pedicels per ploidy, and from 5 leaf petioles per ploidy. Transverse fresh sections between 70-100μm thickness of the petioles and stems were obtained with a rotary microtome equipped with a steel knife, then stained with a solution of 1% phloroglucinol in 20% aqueous HCl, which labels lignified tissues (Jensen, 1962), or with calcofluor white for cellulose (Hughes & McCully, 1975). Flower pedicels were fixed in a solution of paraformaldehyde:glutaraldehyde 3:1v/v, then washed, dehydrated with an increasing gradient of ethanol concentrations (10% to 100%), and finally embedded in Technovit 8100 resin. They were then sectioned at 2μm with a Leica DM ultramicrotome and observed through polarized light, which highlights the lignified cell walls of the tracheids, using a Leica DM LB2 microscope.

### Xylem hydraulic conductance in aerial organs of polyploid atemoyas

With the anatomical data, we estimated xylem hydraulic conductance across ploidies by applying the Darcy’
ss law (Darcy, 1856), which appears as a good model that explains differences in bulk flow (Q: m^3^ s^-1^) through a porous material, taking into account the pressure between two different extremes and considering the area of the porous material, and its conductivity, as to:

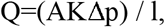

where *A* is the cross-sectional area (m^2^) of the xylem and *K* the specific hydraulic conductivity (m^2^ Pa^-1^ s^-1^), *Δp* is the pressure differential between the soil and the leaf (MPa) under good irrigation, and *l* the distance (m). The total vessel number per cross sectional area was quantified and included as to:

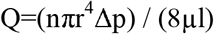

Thus, K= (nπr^4^) / (8μA), where *n* is the average number of vessels, *r* is the radius of individual vessels (m), *μ* the sap viscosity (1 mPa^-1^), and *A* the average xylem area (m^2^). It has to be noted that this calculation gives an overestimation of xylem conductivity, because it does not take into account the distribution of vessel sizes or the resistance of the end walls (connections between conduits) (Roth-Nebelsick *et al*., 2021; Couvreur *et al*., 2018). In addition, the possible contribution of tracheids, with a much smaller diameter, was assumed as minimal compared with vessels. Nevertheless, this method gives a clear prediction of the influence of anatomical traits on the variation of flow rate through the xylem across ploidies.

### Grafted atemoya polyploids under controlled environmental conditions

Due to the difficulty to control environmental conditions in the field, and the long time needed to obtain adult plants from seeds, we decided to make replications of the material maintained in the field by grafting scions from three adult plants, one from each ploidy, onto diploid *A. cherimola* seedling rootstocks during the spring of 2017. As far as we are aware, no interploid grafting had been done before in this species, and the approach worked successfully for 15 scions per ploidy (n=45). After one year of plant acclimation in the greenhouse, the ploidy of the grafted plants was confirmed by flow cytometry as described above. In 2018, 30 grafted plants were transferred to larger ∼0.5 m^3^ Smartpot fabric containers (Root Control, Inc., USA), with 200kg of a 50:50 bark:promix soil, and the surface of the soil was covered with the same fabric to avoid excessive evaporation. While these plants were used for the irrigation treatments (see below), the rest of the plants (15) were kept in well-irrigated-smaller pots in the greenhouse, for an easier transportation to the lab for experiments with dye perfusion.

### *In vivo* xylem sap flow speed in leaves and flowers of grafted polyploids

To monitor the speed of the xylem sap flow *in vivo*, we used nine well-irrigated grafted trees, three per ploidy. A 5mg mL^-1^ esculin hydrate fluorescent dye dissolved in 85% aqueous acetonitrile was perfused manually in the morning of different days (11:00am approximately), with a surgical syringe with a 0.26mm diameter needle in the vascular tissues of the petiole (noted by the stiffness) of three leaves per ploidy and of the flower pedicel of three flowers per ploidy. The room had natural light, but the treated leaves were partially shaded to enhance contrast of the signal provided by an external fluorescent Nightsea UV light. Time-lapse images were recorded every 10s for at least 25 min, using a LeicaS6D dissecting scope connected with a Leica MC190HD camera and the LAS X software. While shading might induce closing of the stomata, the light in the room was enough to keep stomata open. Indeed, we carefully observed the dissemination of the dye through the stomata as a direct evidence of the dye running through the xylem pathway (Video S1). The TIFF images obtained (between 150 and 300 per sample) had a resolution of 10.99μm/pixel (leaves), and 2.10 μm/pixel (carpels), and were segmented with the Image J software based on the default color thresholding method, using an HSB model for color space selection. Threshold values ranged from 116 to 184 and fell into the blue range of the channel, while saturation and brightness were adjusted manually to refine the signal. For each image stack, and given that fluorescence was obscured in the major veins due to the tissues that surround the xylem, a secondary vein was selected in leaves as a region of interest (ROI), its thickness measured, and pixel accumulation with time computed following the Stack Profile Data plugin version 1.0 (24-Sept-2010, Michael Schid). Then, the velocity (V) of the dye was calculated as to:

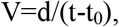

where *d* is the distance (μm), *t* the time (s) of the signal at the end of the ROI, *t*_*0*_ the time (s) at which the signal started to be observed in the ROI.

In the flowers, the exposed carpel surface was the ROI, the accumulation of pixels was computed with the Analyze particles plugin, and the velocity of dye accumulated was calculated on a per area basis (because it was not directional) to:

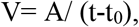

where *A* is the area (μm^2^) of each carpel at the maximum accumulation of pixels; *t* the time (s) at the maximum accumulation of pixels; *t*_*0*_, the time (s) at which the first pixel was observed.

### Irrigation treatments in grafted atemoya polyploids

Our final goal was to test differences in the physiological behaviour of grafted atemoya polyploids under different irrigation intensities and, thus, we severely reduced water applied to the soil compared with the irrigation applied to trees grown in the field. In 2018, we installed drippers that applied water once per week in four pulses of 15 min (temporal separation to avoid the creation of water bulbs in the soil), with three irrigation volumes in three plants per ploidy: good irrigation (16L), medium irrigation (8L), and low irrigation (4L). Soil matric potential was monitored daily using two tensiometers positioned at 15cm and 45cm depth in one pot per ploidy and irrigation regime. To correct for soil evaporative losses, the values obtained from one pot per treatment with no plants but with the tensiometer were subtracted from the pots with plants for each day of measurement. Averages of corrected values were obtained per month (kPa) and transformed to MPa.

### Aboveground organ morphology and gas exchange variables in grafted atemoya polyploids under different irrigation treatments

One year after greenhouse acclimation, we first tested whether scions maintained the characters associated with ploidy compared with the original seedlings grown in the field. Two branches per plant (n=60), were used to quantify branch length/width, leaf number, node density, flower size (length to width ratio), and flower density. Plant phenology was further monitored under greenhouse conditions every other week along the growing season (April to June) by recording the number of nodes bursting, counting the number of leaves in two branches per ploidy and treatment (n=60), and by measuring leaf expansion (length from the lamina insertion into the petiole to the tip of the leaf) in two coetaneous leaves per plant and ploidy (n=60).

To test the effects of ploidy and irrigation on photosynthetic performance, gas exchange measurements were taken for two consecutive days in two fully expanded leaves per plant (n=64), and two days after the irrigation event. First, we searched for possible differences between morning and midday, in 2019, performing initial measurements at both early morning (9:30-11:00 h), and midday (12:30-14:00 h). Since no differences were found between the two measurements, only one measurement was taken at midday in July 2020, when transpiration was at the maximum peak under the subtropical climatic conditions of the greenhouse (22°C, RH=50%, VPD <1.2 kPa, and saturating photosynthetically active photon flux density (PPFD) 800 μmol quanta m^−2^s^−1^). We used an open gas exchange system Li-6400 (LICOR Inc., USA) equipped with a LED-light source (6400-02B) and with a CO_2_ mixer (6400–01). Flow rate was 500 mL min^−1^, and the reference CO_2_ was 400 μmol CO_2_ mol^−1^. Net CO_2_ assimilation rates (A_N_), stomatal conductance (g_s_), intercellular CO_2_ concentration (C_i_) and transpiration rates (E) were estimated with established equations (Farquhar *et al*., 1980). Furthermore, intrinsic water use efficiency (A_*N*_*/*g_*s*_), instantaneous water use efficiency (A_*N*_*/*E), and the ratio of assimilation per internal carbon in the leaves (A_*N*_*/*C_i_) were calculated. These values were also compared with similar measurements of the original plants in the field, and no significant differences were obtained. Concomitantly, the predawn and midday water potential of the leaves (Ψ_l_) were measured during the same days of gas exchange analysis in two leaves per plant (n=64), using a Scholander chamber (Scholander *et al*., 1965) model 600 (PMS Instrument, Corvallis, Oregon, USA).

### Plant hydraulic conductance (K_plant_) of atemoya polyploids

Using the data of soil matric potential for the specific day of gas exchange measurements, as well as leaf water potential in the same plant, and the evapotranspiration values, all measured in the greenhouse, we calculated the whole plant hydraulic conductance using the Darcy’
ss law (Darcy, 1856; Brodribb *et al*., 2021), but with the evaporative flux method under steady state conditions, considering both minimal branching of the young grafted trees, and a lack of internal capacitances, comparing across ploidies and irrigation conditions. We used the formula:

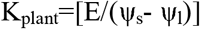

where *K*_*plant*_ is the conductivity of the whole plant (Kg m^-2^ s^-1^ MPa^-1^), *E* the maximum evaporative flux (mmol H_2_O m^-2^ s^-1^), Ψ_*s*_ the soil matric potential (MPa), Ψ_*l*_ the minimum water potential of the leaves at midday (MPa). *K*_*plant*_ was calculated for each plant in particular, and the resulting values averaged.

### Statistical analyses

Anatomical data (number and dimensions of vessels, and xylem areas) of petioles, pedicels and stem of comparable diameters were analysed with a one way ANOVA at a *p*<0.05.

With the results of dye velocity in at least three leaves per ploidy, we applied generalized linear models taking velocity as a dependent variable and vein thickness, ploidy and their intersection as independent variables at a *p*<0.05. Then, we used ANOVA to compare velocity in veins of similar thickness across ploidies, and for the accumulation rates of the dye in carpels (slopes of the ascending pixel cumulative curves). Same models were applied to the phenology measurements of leaf number and size taking as independent variables ploidy and irrigation treatment.

The effects of water treatments and ploidy level on the gas exchange variables and physiological parameters of the *atemoya* trees were evaluated by a two way ANOVA that took as independent variables either ploidy and irrigation, or K_plant_ and irrigation, using IBM SPSS Statistics 2019 v26 analytical software. Normality and homogeneity assumptions were tested prior to ANOVA, using the Kolmogorov-Smirnov and Levene’s tests, respectively. Significant differences were considered at the 5% probability level unless otherwise stated. When significant differences were observed, Duncan’s multiple range test (MRT) was used to compare the mean values.

## Acknowledgements

We are grateful to Jorge Lora and Ruth Aranda Nebot for help with greenhouse experiments, José Carlos Cabrillana for field images, and the technical staff of the IHSM-CSIC-UMA for help with experimental setup. We are also grateful to Miguel Barceló-Anguiano and José Antonio Saavedra for their help with microscopy and field analysis. JML was financed by a ComFuturo Project from the Fundación General CSIC (FGCSIC), and by two RTI Projects (100-900-0000/AEI/10.13039/501100011033 and PID2021-129074OB-I00) from the Agencia Estatal de Investigación-Ministerio de Ciencia e Innovación, Spain, and a LINKB20067 from CSIC. JIH was supported by PID2019-109566RB-I00 funded by MCIN/AEI and ERDF A way to make Europe (MCIN/AEI/10.13039/501100011033).

## Author contributions

J.M.L. designed, supervised the work, and wrote the manuscript. N.B.M. performed in vivo flow velocity trials and statistical analysis. A.F. performed anatomical evaluations and processing of data. E.M.F. performed field evaluations and statistics. J.I.H. supervised the work and the manuscript.

## Data Availability

The data that support the findings of this study are available from the corresponding author upon reasonable request.

## Supporting Information

Fig. S1 Correlation between branch length and diameter for diploid (purple), triploid (brown), and tetraploid (blue) trees grown on their own roots in the field.

Fig. S2 Leaf phenology of *Annona cherimola* x *A. squamosa* diploid (pink), triploid (brown), and tetraploid (blue) genotypes grown in the greenhouse.

Fig. S3 Relative humidity (blue) and temperature (red) in the greenhouse during 2019 and 2020.

Fig. S4. Total leaf number per branch in diploid (pink), triploid (brown), and tetraploid (blue) atemoya scions at the end of the growth season (June 2020), as a function of treatment (x axis).

Fig. S5. Stomatal conductance of *Annona cherimola* x *A. squamosa* diploid (pink), triploid (brown), and tetraploid (blue) genotypes grown in the greenhouse under three irrigation regimes.

Table S1 Gas exchange variables.

