## Supplementary materials for "Hydraulic tradeoffs underlie enhanced performance of polyploid trees under soil water scarcity"

**Article title:**

**SUPPORTING INFORMATION TITLES AND LEGENDS**

**Fig. S1 Correlation between branch length and diameter for diploid (purple), triploid (brown), and tetraploid (blue) *Annona cherimola* x *A. squamosa* trees grown on their own roots in the field.**

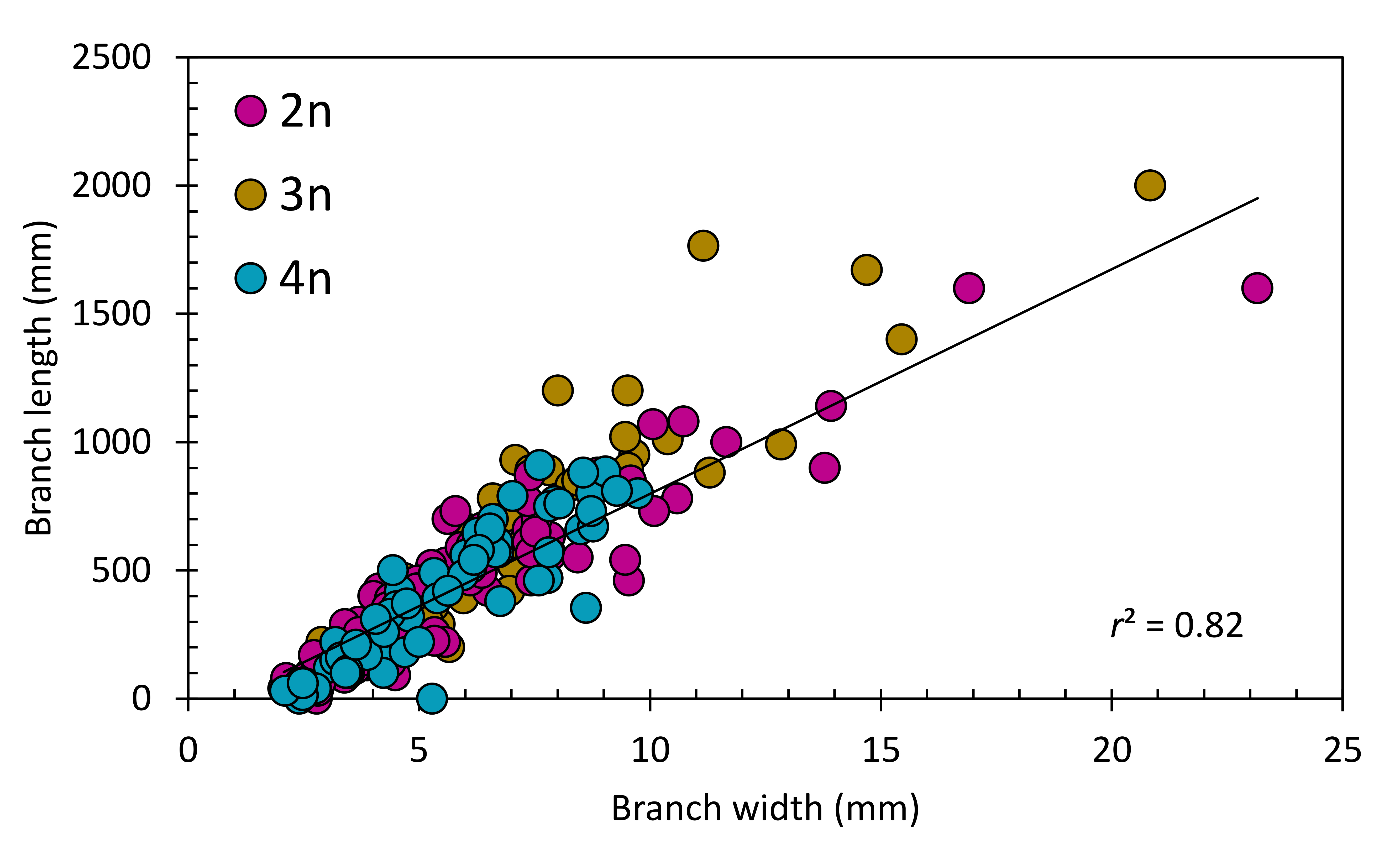

**Fig. S2 Leaf phenology of *Annona cherimola* x *A. squamosa* diploid (pink), triploid (brown), and tetraploid (blue) genotypes grown in the greenhouse. (a)** Monitoring total leaf number per branch with time in the growing season of 2020. **(b)** Leaf elongation rates during the growing season of 2020; inset shows the mature leaf areas for all three ploidies in the greenhouse. Bars represent standard error, and letters show significant differences after ANOVA test, both at *p*<0.05. ­­­
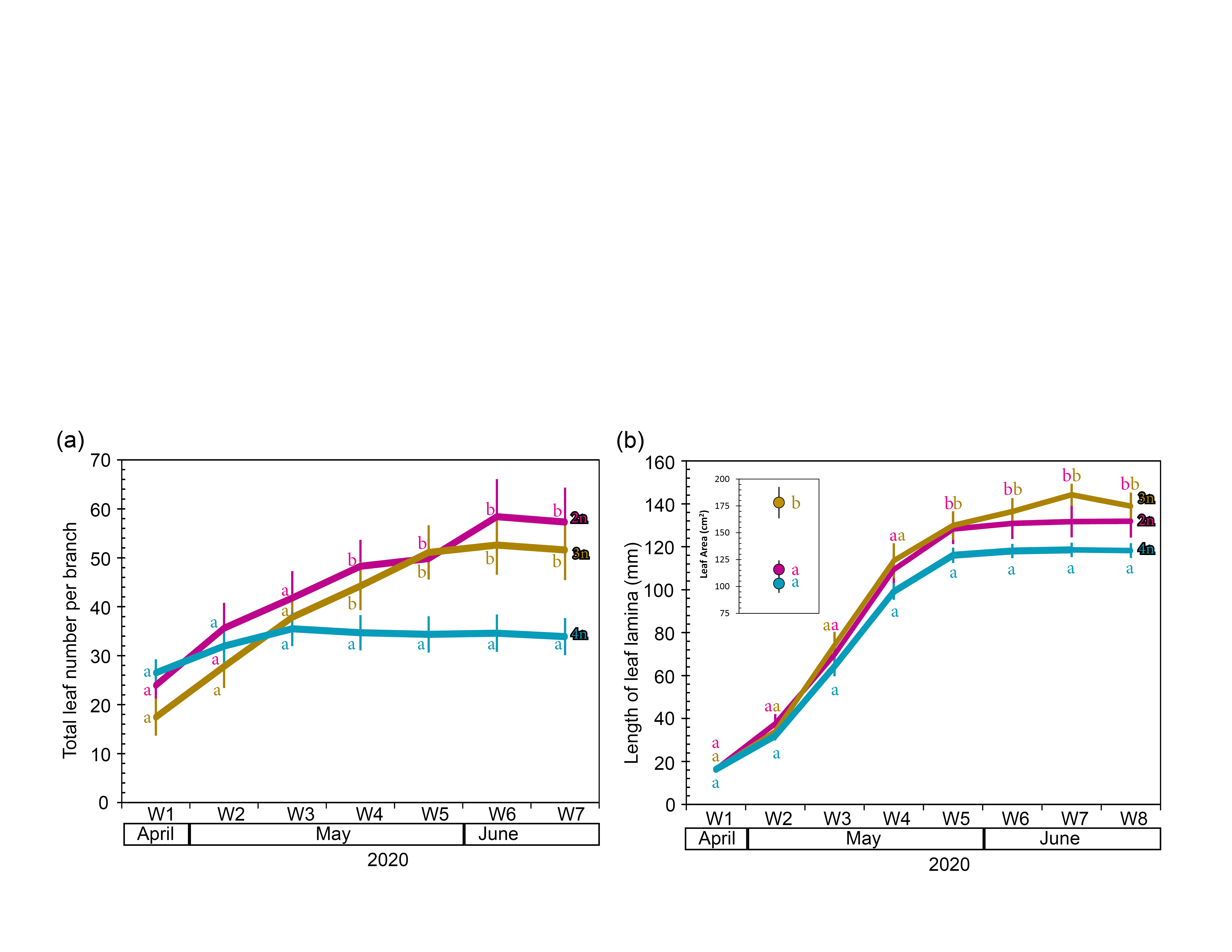

**Fig. S3** **Relative humidity (blue) and temperature (red) in the greenhouse during 2019 and 2020.** Colors represent daily fluctuations between maximum daily averages on top and the minimum night averages at the bottom of each plot. Shaded months represent the dates of the growing period.

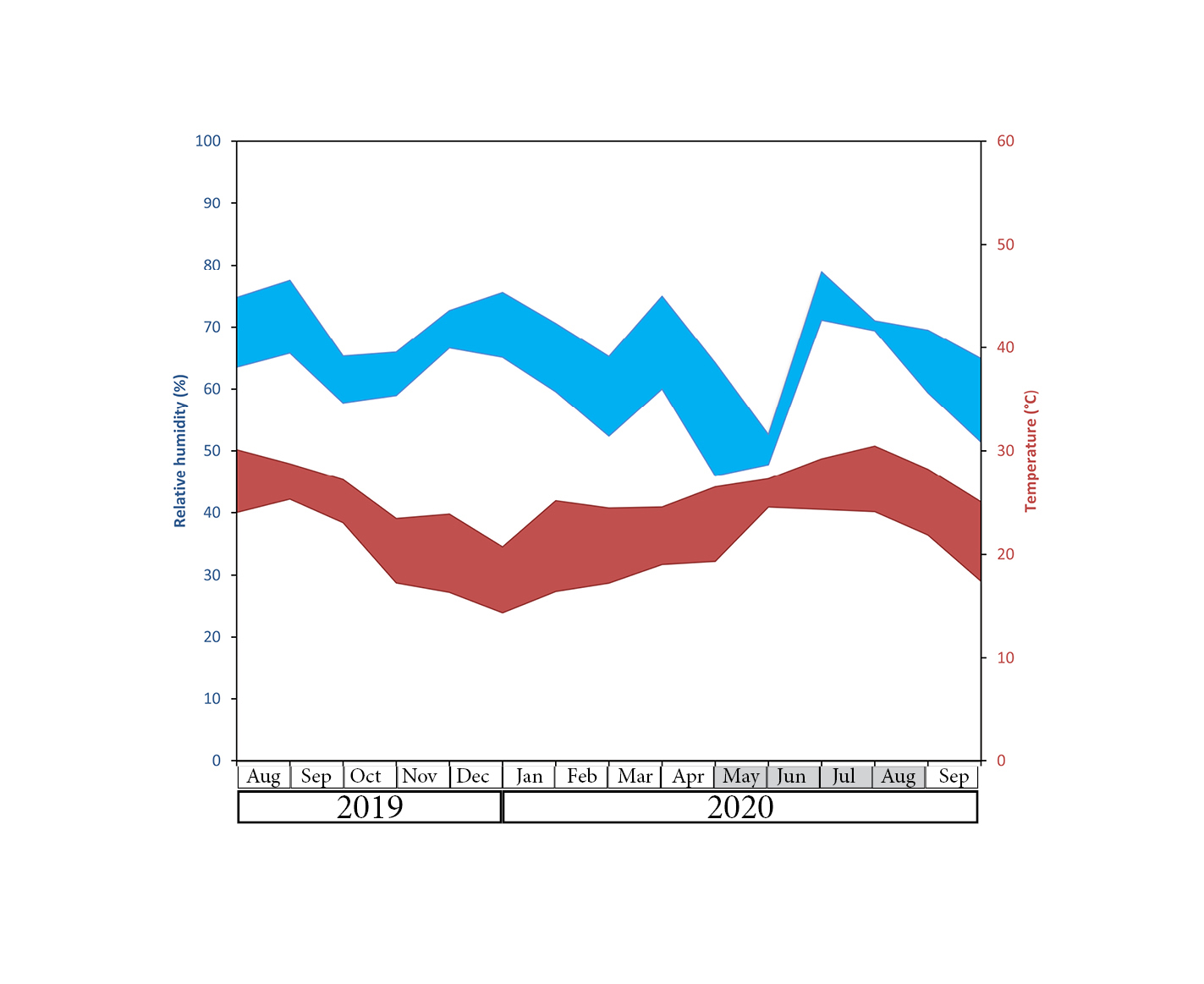

**Fig. S4. Total leaf number per branch in diploid (pink), triploid (brown), and tetraploid (blue) *Annona cherimola* x *A. squamosa* scions at the end of the growth season (June 2020), as a function of treatment (x axis).** Bars represent standard error at *p* < 0.05.

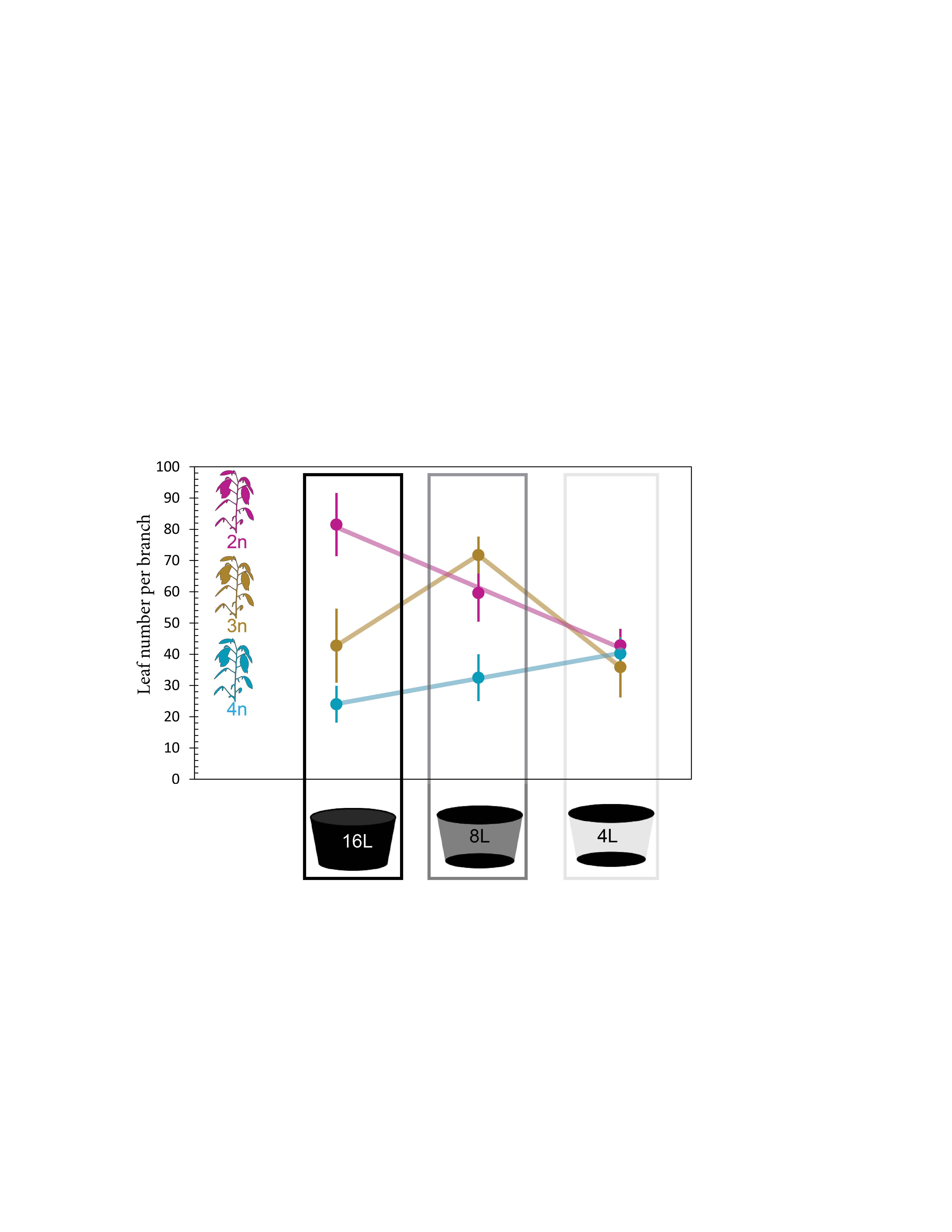

**Fig. S5. Stomatal conductance of *Annona cherimola* x *A. squamosa* diploid (pink), triploid (brown) and tetraploid (blue) genotypes grown in the greenhouse. Letters indicate significant differences after a two-way ANOVA at *p* < 0.05.**

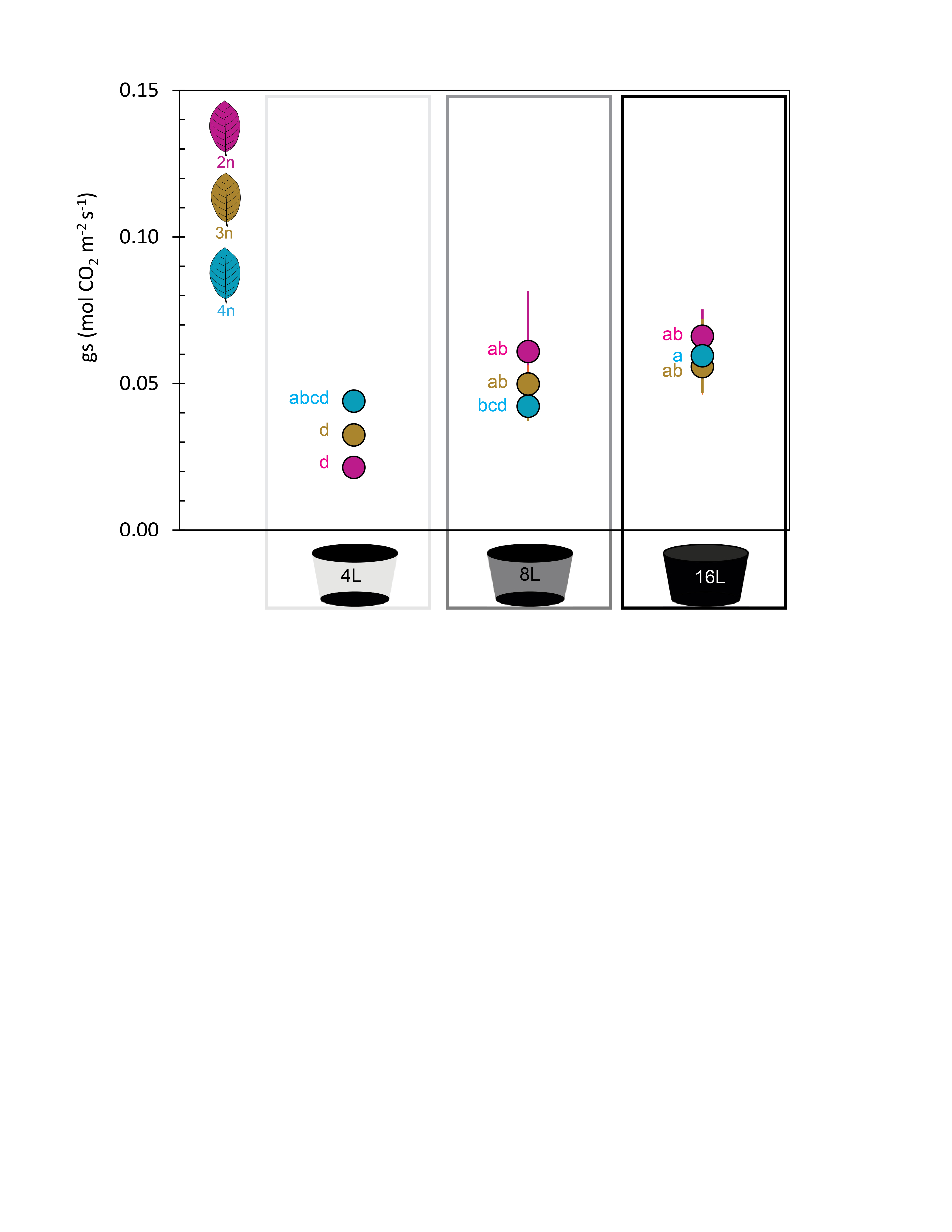

**Table S1. Gas exchange variables.** Net CO_2_ assimilation rate (*A_N_*), stomatal conductance (*gs*), transpiration rate (*E*), intercellular CO_2_ concentration (*Ci*), intrinsic water use efficiency (*A_N_/gs*) and instantaneous water use efficiency (*A_N_/E*) of genotypes of different ploidies of *Annona cherimola* x *A. squamosa* measured in full-water treated plants (FW, 16L), well-watered plants (WW, 8L) and water stressed plants (WS, 4L) in July of two consecutive years (2019 and 2020), except FW, with measurements only for 2020. Each value represents the mean ±SE. Within each variable, different letters indicate significant differences among water treatments and ploidies (*P*<0.05) in 2019 (capital letters) and in 2020 (lower cases).

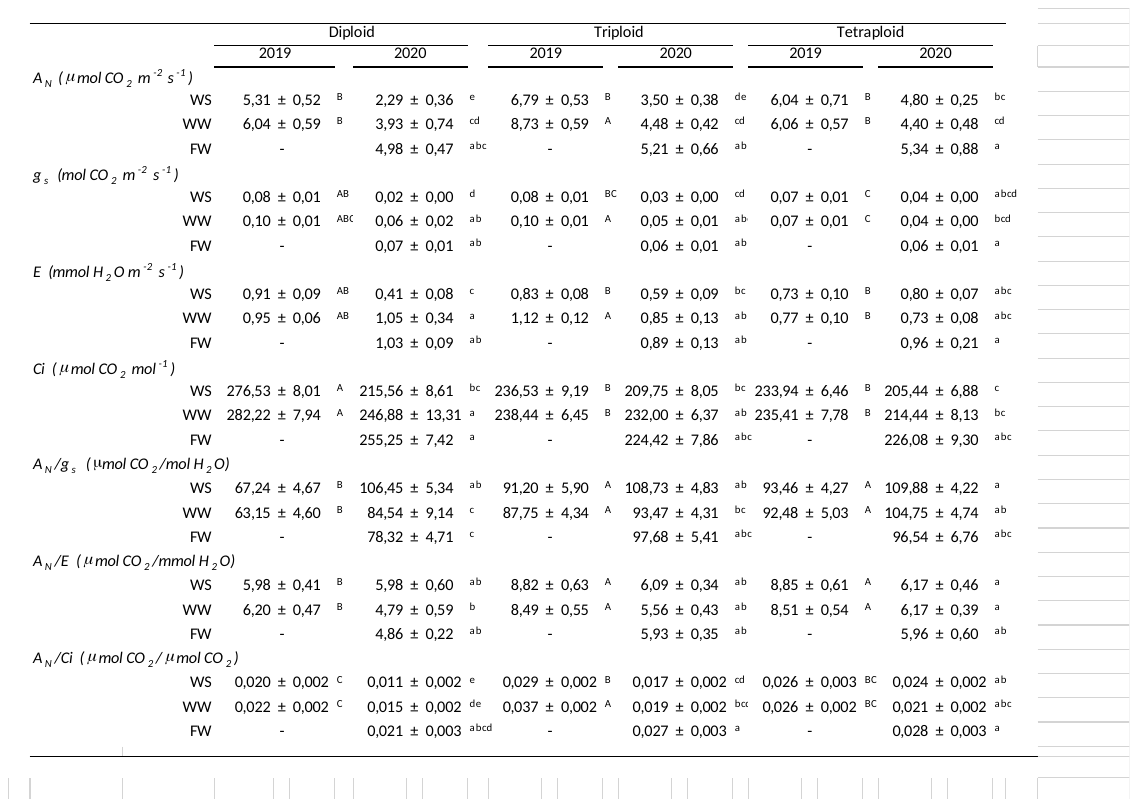
